# Lessons from the analysis of TAD boundary deletions in normal population

**DOI:** 10.1101/2020.04.01.021188

**Authors:** Thomas Smol, Jérôme Sigé, Caroline Thuillier, Frédéric Frénois, Perrine Brunelle, Mélanie Rama, Catherine Roche-Lestienne, Sylvie Manouvrier-Hanu, Florence Petit, Jamal Ghoumid

## Abstract

Topologically Associating Domains (TAD)-boundaries induce spatial constraints, allowing interaction between regulatory elements and promoters only within their TAD. Their disruption could lead to disease, through gene-expression deregulation. This mechanism has been shown in only a relatively low number of diseases and a relatively low proportion of patients, raising the possibility of TAD boundary disruption without phenotypical consequence. We investigated, therefore, the occurrence of TAD boundaries disruption in the general population. Coordinates of 307,430 benign deletions from public databases were crossed with 36 Hi-C datasets. Differences in gene content and gene localization were compared in the TADs, according to the possible disruption of their boundaries by a deletion found in the general population. TADs with no deletion encompassing their boundaries (R-TAD) represented 38% of TADs. Enrichment in OMIM genes as well as in morbid genes was observed in R-TADs and genes in R-TADs were found to localize closer to the boundaries. Our results support recent publications tempering the impact of breaking TADs on gene expression with a majority of broken TADs in the general population. A subgroup of R-TAD emerges from this analysis with enrichment in disease genes and their coordinates could be used to annotate CNV from pangenomic approaches to enhance data interpretation.

## Introduction

Development of Hi-C has greatly contributed to elucidate 3D genome organization. The technology provided information on genome-wide interactions within a single experiment. At a resolution below 40 kb, Hi-C experiments identify chromatin domains, corresponding to Topologically Associating Domains (TADs). These structures are characterized by high level of intra-domain chromatin interactions contrasting with low level of inter-domain chromatin interactions (Szabo et al. 2019). On Hi-C heat maps, TADs appear as pyramid-shaped structures, arising from chromatin-contact enrichment and separated by chromatin-contact depletion zones, referred as TAD-boundaries (Chang et al. 2020). The boundaries of TADs are specific structures playing the role of insulators, *i.e*. preventing chromatin inter-domain interactions. They show not only enrichment in binding sites for DNA-binding proteins, such as CTCF, ZNF143, SMC, and YY1 but also specific histone epigenetic modifications, such as H3K36 trimethylation (H3K36me3), and Transcription Starting Sites (TSS) of housekeeping genes (Hong and Kim 2017).

CTCF-binding sites are the key features of TAD-boundaries. The protein promotes interactions between distant genomic elements by mediating chromatin loop formation. A model of TAD formation has been therefore proposed by loop extrusion: cohesin protein complex progresses along the chromatin, forming a growing loop until it interacts with two CTCF molecules bound with convergent orientation.(Sikorska and Sexton 2020).

TAD-boundaries induce spatial constraints, enabling interaction between regulatory elements and promoters only within a TAD, and preventing improper action of extra-TAD enhancers on these genes. As a consequence, during cell differentiation, genes located within the same TAD are usually coregulated (Szabo et al. 2019).

TAD-boundary disruption has been shown to cause genetic diseases, through a gene-expression deregulation mechanism. First examples of this pathophysiological mechanism were limb malformations due to chromosomal rearrangements of the *WNT6-IHH-EPHA4-PAX3 locus*. For example, while *EPHA4* is not involved in limb development, deletions encompassing this gene result in brachydactyly. Deletions disrupting the boundary between TADs containing *EPHA4* and *PAX3* abolish its insulator-effect and allow improper regulation of *PAX3* by a cluster of limb-specific enhancers from the adjacent TAD (Lupiáñez et al. 2015a). Many others examples of this mechanism have been reported, including TADs containing *LMNB1* or *SOX9* genes (Nmezi et al. 2019)(Franke et al. 2016).

Here we studied TAD boundary deletions in control databases, to investigate the possibility and the occurrence of TAD boundary disruptions in the general population. We found that up to 62% of TADs have a disrupted boundary in the general population, highlighting the possibility of TAD disruption without observable consequence.

## Methods

### CNV collection

Genomic coordinates of benign and likely benign copy number deletions were extracted from ClinVar(Landrum et al. 2016), Database of Genomic Variants (DGV)(MacDonald et al. 2014), DECIPHER (Database of genomic variation and Phenotype in Humans using Ensembl Resources)(Firth et al. 2009), genome aggregation database (GnomAD)(Karczewski et al. 2019) and from control cases described in a previously published series(Coe et al. 2014) (supplemental material table 1). The GRCh37 reference was used for coordinate descriptions. A BED file combining all the 307,430 benign or likely benign copy number deletions was compiled (supplemental material table 1) (Kent et al. 2002).

### Hi-C datasets

Thirty-Six Hi-C datasets available online were used. Seven out of the 36 datasets were generated by Arrowhead method, concerning the cell lines: GM12878, HMEC, HUVEC, IMR90, K562, KBM7 and NHEK (Rao et al. 2014). Twenty-One datasets were generated by the directionality index method applied, namely 5 human embryonic stem-cell derived lineages: H1-ESC, H1-MES, H1-MSC, H1-NPC, H1-TRO; 4 set of human tissues: Aorta-STL002, Liver-STL011, Thymus-STL001 and VentriculeLeft-STL003; and 12 set of human cell lines: A549, Caki2, G401, LNCaP, NCIH460, PANC1, RPMI7951, SJCRH30, SKMEL5, SKNDZ, SKNMC, T470 (Leung et al. 2015; Dixon et al. 2018, 2015). Nine were generated by high-resolution 3D maps from donor tissues: Cortex-DLPFC, Adrenal Gland, Bladder, Bowel Small, Lung, Muscle Psoas, Pancreas, Spleen and Ventricle Right (Won et al. 2016). The GRCh37 reference was used for coordinate descriptions.

### Genomic annotations

Genomic elements were collected from the University of California Santa Cruz Genome Browser database (UCSC)(Rosenbloom et al. 2015) which were generated for most of them by the Encyclopedia of DNA Elements (ENCODE) (ENCODE Project Consortium 2012). Coordinates of 31,842 known canonic isoforms, mapping to Ensembl, were extracted from UCSC TableBrowser (supplemental material table 3) and annotated with OMIM database (n=9,848 genes) (Amberger et al. 2015). A list of genes associated with human diseases was defined from the manually curated HGMD database with a selection of genes with disease-causing variations (n=7,014 genes) (Cardoso-Moreira et al. 2019). Developmental genes were previously defined from mammalian resource on organ development with an analysis of 297 RNA-seq human libraries (n=6,646) (Cardoso-Moreira et al. 2019). Coordinates of enhancers were also collected from GeneHancer database (Fishilevich et al. 2017).

### Crossing data

Annotation of the Hi-C datasets and analyses of overlap between genomic regions were performed with the BEDtools suite (Quinlan and Hall 2010). Four different subgroups of TAD were defined from Hi-C datasets, according to the involvement of their boundaries in a benign and likely benign copy number deletion. Thus, “robust TAD” (R-TAD) corresponds to TADs with no boundary encompassed by one of these deletions; “un-robust TAD” (U-TAD) corresponds to TADs with both boundaries possibly encompassed by one of these deletions; S1-TAD corresponds to TADs with start-boundary possibly encompassed by one of these deletions; and finally, S2-TAD corresponds to TADs with end-boundary possibly encompassed by one of these deletions. We compared gene densities in the different sub-groups of TADs. Gene density was calculated by the ratio of the Open Reading Frame (ORF) length of coding genes within a TAD on the total length of the TAD. This gene density ranged from 0.0, meaning 0% of the TAD length corresponded to a gene ORF, to 1.0 meaning that 100% of the TAD length correspond to gene ORFs. Then, we compared the global gene density (n=31,852 isoforms) and the OMIM genes (n=9,848), developmental genes (DEV genes; n=6,646) and morbid genes (DM genes; n=7,014) densities between R-TAD and U-TAD.

We compared the pLI of genes involved either in R-TAD or in U-TAD. The loss-of-function intolerant metric score (pLI) has been developed on the basis of GnomAD and ExAC databases, and indicates the probability for a gene to be intolerant to loss-of-function variation and to fall into the haplo-insufficiency category (Karczewski et al. 2019).

The intra-TAD organization, *i.e*. gene-TAD boundary distance and intra-TAD enhancer distribution was studied after normalization of all TADs size to 1,000,000bp. The positions of each gene and enhancer were analyzed within all TADs.

### Statistical analysis

The Statistical analysis was performed using the computing environment R 3.6.1 with additional software packages taken from the Bioconductor project(Gentleman et al. 2004): karyoploteR, BSgenome.Hsapiens.UCSC.hg19 and GenomicRanges. Figures were made using the ggpubr and the karyoploteR packages (Gel and Serra 2017). The level of statistical significance was set at 0.05. The reported results were rounded to two decimals.

## Results

### Frequency of TAD boundary disruptions in the general population

Extraction of genomic coordinates of 5 databases provided a collection of 307,430 deletions which were crossed with TAD coordinates from 36 Hi-C datasets. First of all, only a median frequency of 0.38 [0.33 – 0.41] of TAD was considered as robust TADs (R-TAD) (*p<2e−16*) (Figure 1A). The frequency of R-TAD was very similar between all Hi-C datasets with the lowest and highest frequencies observed respectively in Thymus_STL001_Leung2015(Leung et al. 2015) with 0.33 of R-TAD (322 R-TAD out the 984 identified) and in HMEC_Lieberman dataset(Rao et al. 2014) with 0.41 of R-TAD (1,314 R-TADs out the 3,235). The S1-TAD and S2-TAD subgroups frequency is 0.41 [0.37 – 0.44] and unrobust-TAD (U-TAD) frequency is 0.21 [0.19 – 0.25]. A statistical difference was found between the median size of R-TAD and U-TAD, with respective values of 920,000bp [350,000 – 2,360,000] and 1,075,000bp [375,000 – 2,650,000] (*p<2e-16*) (Figure 1B).

**Figure 1:**
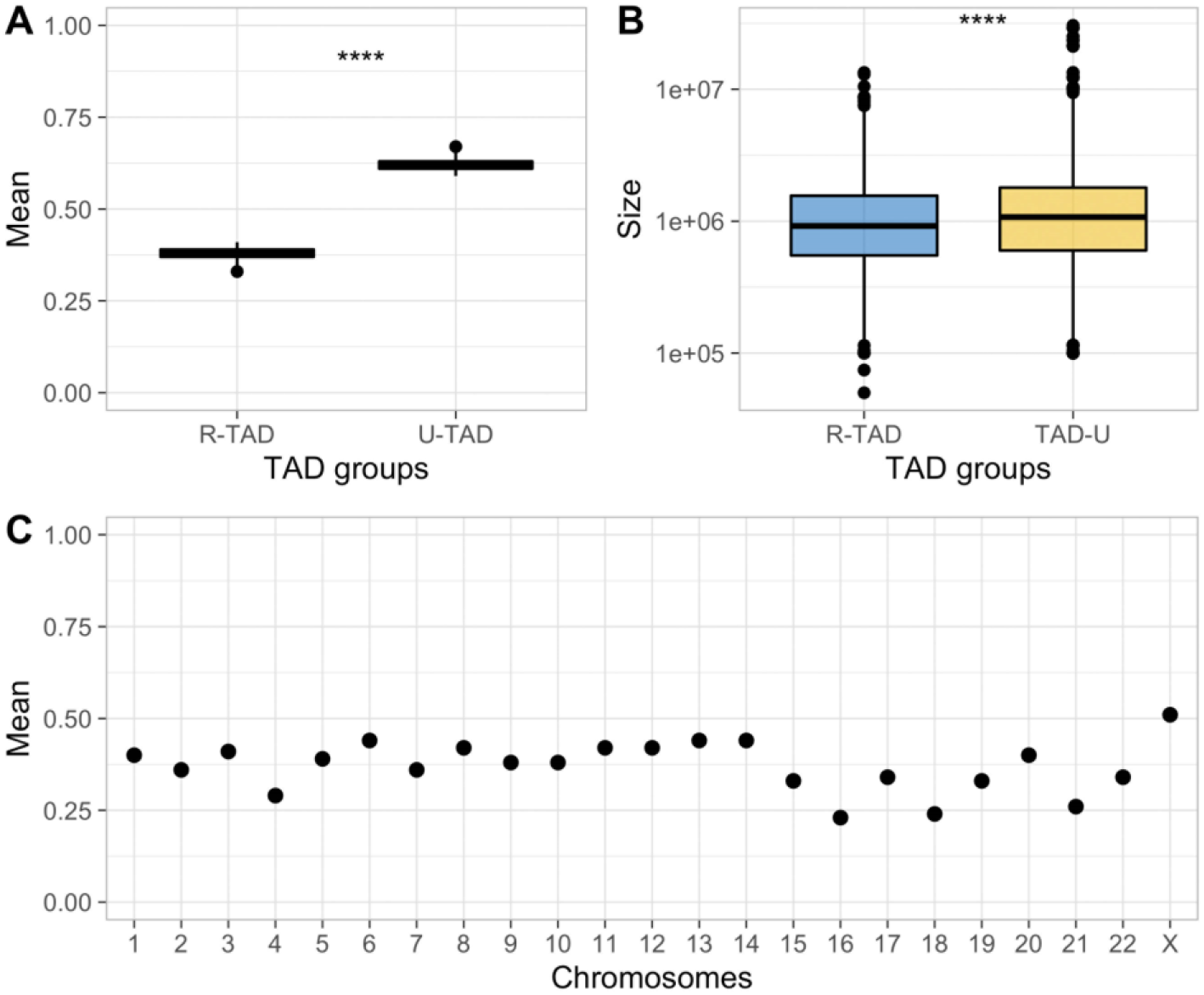
TAD robustness among 36 Hi-C datasets. **(A)** Boxplots of TAD robustness rom 36 Hi-C datasets. The highest level of robustness as determined for HMEC cell line with 0.41 and the lowest for 0.33 for Thymys_STL001_Leung2015. The median robustness as 0.38. Center lines represent the median; box limits are the upper and the lower quartiles and points are outliers. **(B)** Density plots of R-TAD and U-TAD sizes indicate a larger size for U-TAD than R-TAD with respective mean sizes of 1,075,000bp [375,000 – 2,650,000] and 920,000bp [350,000 – 2,360,000] (*p<2e-16*). R-TAD are plotted in blue and U-TAD in yellow, mean values are represented by black bars. **(C)** Percentage of R-TAD according to chromosome.

The repartition of R-TADs was heterogeneous among chromosomes and three different groups of chromosomes could be observed (Figure 1C). The first group, defined by low R-TAD density, comprised chromosomes 16, 18 and 21, with a median R-TAD density of only 0.24 [0.23 – 0.26]. The second group, defined by medium R-TAD density, corresponded to the remaining autosomal chromosomes and was associated with a median of 0.39 [0.29-0.44]. In this group, median density ranged from 0.29 in chromosome 4, to 0.44 in chromosomes 6, 13 and 14. The third group, defined by high TAD-robustness, was only represented by chromosome X with R-TAD median of 0.51 (Figure 1C). The difference of R-TAD repartition between these three groups was found statistically significant (*p<2e-16*). The distribution of R-TAD and U-TAD was not likely to be associated neither to chromosomal banding organization, nor to centromere or telomere areas (Supplemental Figure 1).

### Gene enrichment in the different TAD sub-groups

Comparison of global gene density, between R-TAD and U-TAD showed no difference between the two groups with a median of 0.52 and 0.51 respectively (*p=0.43*) (Figure 2A). A higher OMIM genes density(Amberger et al. 2015) was observed in R-TADs with 0.27 compared to 0.19 in U-TADs (*p=0.003*) (Figure 2B). Differences in gene density were also observed for DEV-genes and morbid genes between R-TADs and U-TADs with respectively 0.10 *vs* 0.06 (*p=0.03*) and 0.18 *vs* 0.07 (*p=0.0002*) (Figure 2C and 2D). Comparing pLI of genes located either into R-TADs or into U-TADs revealed that genes located in R-TADs presented a statistically higher pLI with a mean value of 0.26 [6.31e-13 – 0.99] than genes in U-TADs with a mean pLI of 0.22 [1.29e-14 – 0.87] (*p<2e-16*) (supplemental Figure 2).

**Figure 2:**
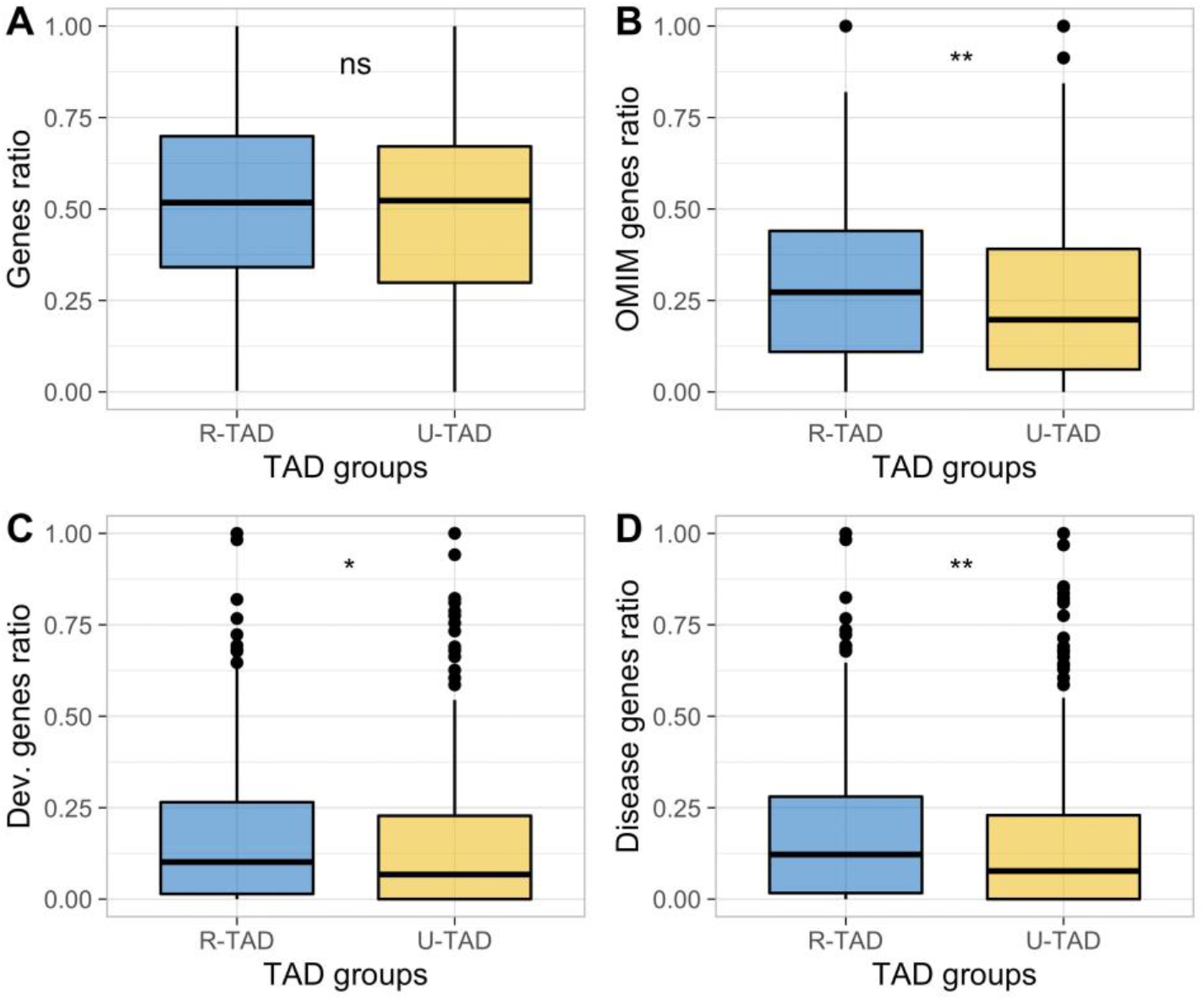
Comparison of gene enrichment between R-TAD and U-TAD. **(A)** Boxplots of genes ratio in R-TAD and U-TAD subgroups. No statistical difference was observed in genes ratio between R-TAD (0.52) and U-TAD (0.51) (*p=0.43*). **(B)** Boxplots of OMIM genes ratio. A higher OMIM genes enrichment was observed in R-TAD with a mean ratio of 0.27 compared with 0.19 in U-TAD (*p=0.003*). **(C)** Boxplots of developmental genes ratio. A higher developmental genes enrichment was observed in R-TAD with a mean ratio of 0.10 compared with 0.06 in U-TAD (*p=0.03*). **(D)** Boxplots of disease genes ratio. A higher disease genes enrichement was observed in R-TAD with a mean ratio of 0.18 compared with 0.07 in U-TAD (*p=0.0002*).

### Distribution of genes and enhancers in the different TAD sub-groups

Global gene and enhancer distributions followed a U-shape organization with a polar enrichment (Figure 3A and 3B). The median distance of genes from the 5’ or 3’-TAD boundaries was estimated respectively to ±199,076bp for R-TADs and ±218,282bp for U-TADs. Thus, R-TAD genes were located 19,206bp closer to boundaries than U-TAD genes (*p<1e-13*).

**Figure 3:**
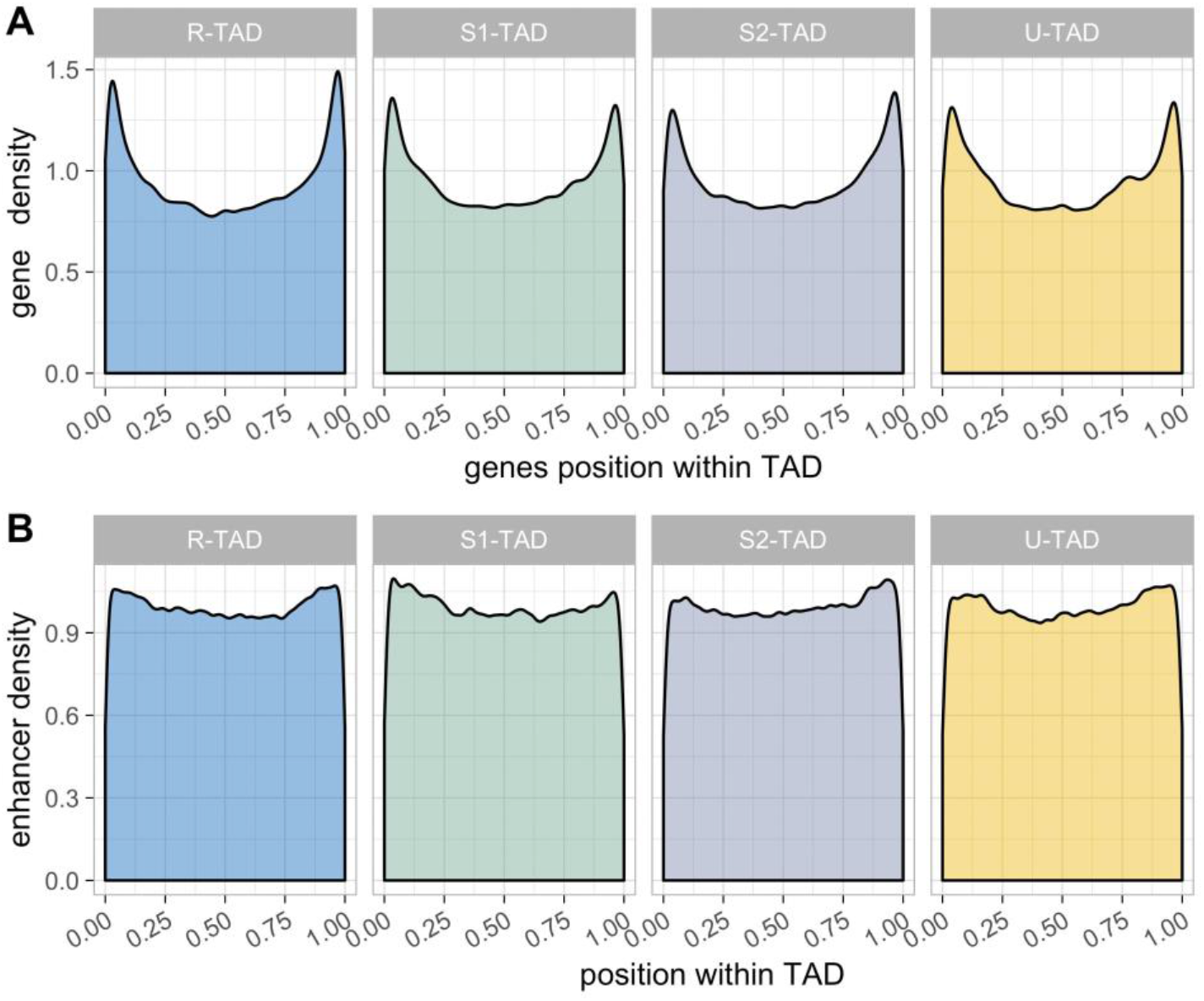
Density plots of genes and enhancers distribution in each TAD sub-group. **(A)** Genes distribution within the TADs. Genes are organized in U-shape with a significant higher density close to undisrupted TAD boundaries. (B) Enhancers distribution within the TADs.

The median gene-boundary distances were respectively of ±35,900bp and ±38,080bp for undeleted and deleted boundaries (*p<2e-16*). The median OMIM gene-boundary distances were respectively of ±35,680bp and ±39,400bp for undeleted and deleted boundaries (*p<2e-16*). The median developmental gene-boundary distances were respectively of ±34,140bp and ±39,480bp for undeleted and deleted boundaries (*p<2e-16*). The median disease gene-boundary distances were respectively of ±34,140bp and ±37,946bp for undeleted and deleted boundaries (*p<2e-16*).

Asymmetric TADs, *i.e* TADs with only one boundary encompassed by a deletion, were associated with an asymmetric distribution of genes. The median distance of genes from undeleted and deleted boundaries was estimated respectively to ±201,234bp and ±221,012bp (*p<1e-13*). At deleted TAD boundary loci, a lower gene enrichment was observed (supplemental Figure 3A and 3B).

## Discussion

Development of 3D genome visualization techniques allowed accumulation of knowledge about genome architecture and have led to a higher understanding of genomic regulation. Elucidation of the regulatory role of TAD boundaries allows understanding how particular chromosomal imbalances, distant from genes, could result in gene-expression deregulation and in diseases (Lupiáñez et al. 2015b). TAD-boundary insulator abilities prevent regulatory elements from interacting with gene promoters located outside the TAD. Thus, boundary disruptions abolish this spatial constraint and would allow ectopic enhancer/gene-promoter interactions. Limb malformations caused by chromosomal imbalances at *EHPA4* locus exemplify this mechanism (Lupiáñez et al. 2015a).

Even if TAD-boundary disruption was considered as a well-established model of deleterious mechanism underlying genetic diseases, this mechanism has been shown in only a relatively low number of diseases (Spielmann et al. 2018). In addition, large cohort studies identified TAD-boundary disruption as the disease-causing mechanism only in a low proportion of patients, ranging from 0.3% to 11.8% (Di Gregorio et al. 2017; Ibn-Salem et al. 2014). In light of these data, we wondered whether TAD boundary disruptions could occur without resulting in an observable phenotype. To do this, we assessed the proportion of TAD boundaries encompassed by at least a benign or likely-benign deletion. We consider only deletions, because interpretation of small duplications or inversions are complex and non-consensual. We also extracted TAD boundaries coordinates from 36 HI-C datasets available online, providing TAD-boundary data in different regulatory contexts: immortalized cell lines (GM12878, HMEC, HUVEC, IMR90, K562, KBM7, NHEK, A549, Caki2, G401, LNCaP, NCIH460, PANC1, RPMI7951, SJCRH30, SKMEL5, SKNDZ, SKNMC, T470); human embryonic stem-cells derived lineages (H1-ESC, H1-MES, H1-MSC, H1-NPC, H1-TRO) and human tissues (aorta, liver, thymus and heart left ventricle, cerebral cortex, adrenal gland, bladder, small bowel, lung, psoas muscle, pancreas, spleen and heart right ventricle). Considering TADs in various cellular and tissue contexts was important, since single-cell Hi-C assays showed that TAD structure could be variable in a same tissue (Chang et al. 2020). We collected genomic coordinates of 307,430 benign and likely benign copy number deletions from various public databases (ClinVar, DGV, DECIPHER and Phenotype in Humans using Ensembl Resources, GnomAD and from control cases of previously published series). We crossed the data to determine which TADs had no boundary encompassed by a benign or likely-benign deletion and called them R-TADs (robust-TAD). We determined also TADs with deletion of their 5’-boundary (S1-TAD), 3’-boundary (S2-TAD) or deletion of their two boundaries (U-TAD; unrobust-TAD) reported in the general population.

We found that only 38% of the TADs belong to R-TAD and 21% belong to U-TAD. Most TADs (41%) had either their 5’-boundary (S1-TAD) or their 3’-boundary (S2-TAD) encompassed by a benign or likely benign deletion (Figure 1A). These results highlight that a majority of TAD boundaries could be deleted, resulting in no or minor phenotype.

To assess differences between these four sub-groups, we calculated their relative enrichment in OMIM genes, DEV-genes (developmental genes) and disease-causing genes. We also determined the relative distance between genes and TAD boundaries. We found that global gene density was similar in the 4 sub-groups of TADs (Figure 2). However, density in OMIM genes, DEV-genes and disease-causing genes was higher in R-TADs. In addition, R-TADs were likely to be enriched in genes with a high pLI. The four sub-groups were also different in term of genes and enhancers distribution along TADs (Figure 3). Genes and enhancers localized preferentially at the 5’ - 3’-boundaries of the TADs, resulting in a U-shape distribution (Figure 3A and 3B). The R-TADs showed a higher gene density close to the 5’- or 3’-TAD boundaries. We observed asymmetric distribution of genes in S1-TADs and S2-TADs. Gene density was lower at the 5’-TAD boundaries of S1-TADs and at the 3’-TAD boundaries of S2-TAD (Figure 3). OMIM genes, DEV-genes and disease causing-genes were found to locate preferentially closer to the undeleted boundary compared to other genes. This was consistent with the fact that TAD insulating capacity is relatively weak (Chang et al. 2020). By contrast, deletion of TAD boundaries in individuals without observable phenotype, probably indicate that the expression of genes within U-TADs was less dependent on the insulator abilities of the TAD boundaries, explaining the shape of the genes distribution along U-TADs and the lower enrichment in diseases-causing genes.

A model can emerge from these data. TAD-boundaries are located in *loci* with high enhancer and gene density, where proximity would enable ectopic gene/enhancer interactions. If genes surrounding the boundary are requiring insulation from ectopic enhancer activity (typically OMIM genes, DEV-genes and disease-causing genes), genes would possibly undergo a selection pressure to localize close to the TAD-boundary in order to take full advantage of its insulation abilities. Along the same lines, we can observe that chromosome X shows the highest R-TAD density, when it shows a high OMIM-, disease- and DEV-gene density. This contrasts with chromosomes 16, 18 and 21, showing low gene density and the lowest R-TAD density (Figure 1C). The critical role of the TAD-boundaries for this subset of genes is highlighted by the fact that deletions of these TAD-boundaries are not found in the general population. This situation corresponds to S1-, S2-TADs and R-TADs. TADs comprising *EPHA4* and *PAX3* fit this model. Normal expression of *PAX3* requires prevention from ectopic action of limb specific regulatory elements located within TAD containing *EPHA4*. Deletion of the TAD-boundary between *EPHA4* and *PAX3* result in abnormal expression of *PAX3* and in brachydactyly (Lupiáñez et al. 2015a). We found that TAD-boundary between both genes, regardless of the tissue or cellular context, was not encompassed by any deletion found in the general population. However, the model explains not all the situations. In the case of *LMNB1-* related leukodystrophy, three different deletions upstream the gene have been shown to cause the condition. The minimal critical region required for the disease phenotype, spanning about 167 kb, disrupts the 3’-boundary of TAD containing *LMNB1*. It is assumed that the TAD disruption allows ectopic action of a brain specific regulatory element, called *Enh-B*, on *LMNB1* promoter (Nmezi et al. 2019). Surprisingly, we identified 38 benign or likely-benign deletions encompassing the 3’-boundary of *LMNB1-containing* TAD, using data from the 36 Hi-C assays. All 38 deletions were smaller than the minimal critical region span, the larger deletion being about 10.5 kb. Thus, deletion of the TAD boundary is likely not sufficient to cause leukodystrophy. To ectopically regulate *LMNB1, Enh-B* should probably be placed close to the gene, explaining why small deletions were not associated with the disease. The same mechanism could also be involved in brachydactyly due to *EPHA4* locus deletion. Indeed, size of the deletions causing brachydactyly is 1.75–1.9 Mb, allowing positioning of limb specific regulatory sequences from TAD containing *EPHA4*, in direct proximity to *PAX3* promoter. The enhancer-gene distance is probably also critical to result in disease, when TAD boundary is disrupted. In the case of F-syndrome, an inversion involving the TAD boundary between the *WNT6-IHH* and *EPHA4* TADs results in gene deregulation. The inversion abolishes insulation from limb-specific regulatory elements located within *EPHA4-*containing TAD and places these elements next to *WNT6*. Same phenomenon occurs in case of *WNT6* locus duplication, when limb-specific regulatory elements are placed in direct proximity to the duplicated *WNT6* copy, resulting in polydactyly (Lupiáñez et al. 2015a).

Another evidence that the TAD-boundary disruption by itself is not sufficient to result in disease is the robust expression of *Shh* and *Sox9* despite TAD perturbation. Deletion of CTCF-binding sites at the boundaries of the TAD containing *Shh* has no detectable effect on *Shh* expression patterns or levels during development. Expression deregulation occurs when enhancers within the TAD are deleted, or in case of large deletion of its 5’-boundary (Williamson et al. 2019). In the TAD containing *Sox9*, deletion of all major CTCF-binding sites at the boundaries showed no obvious effects on gene expression. Gene misexpression causing disease phenotypes were found to be achieved by redirecting regulatory elements. Thus, disease-related gene activation is not induced only by loss of insulation but requires CTCF-dependent redirection of enhancer–promoter contacts (Despang et al. 2019).

## Acknowledgements

We gratefully acknowledge the European Reference Network ITHACA for support. We thank all members of the EA7364 RADEME team of Lille University for their comments.

## Author contributions

T.S. and J.G. designed the study. T.S., J.S., JG., F.P. and J.G. analyzed the results. T.S. and J.G. wrote the manuscript. All authors discussed the results and commented on the manuscript.

---

**Supplemental Figure 1: Karyoplot of all chromosomes with distribution of R-TAD and U-TAD.** Robust TAD are depicted in blue, unrobust TAD in yellow. Karyogram with G-banding is shown for all chromosomes.

**Supplemental Figure 2:** GnomAD pLI score of R-TAD genes and U-TAD genes. R-TAD are plotted in blue and U-TAD in yellow. R-TAD presented a statistically higher pLI with a mean value of 0.26 [6.31e-13 – 0.99] than genes in U-TAD with a mean pLI of 0.22 [1.29e-14 – 0.87] (p<2e-16).

**Supplemental Figure S3:** Density plots of genes and enhancers distribution in each TAD subgroup in the 5’ extremity of TAD (A) and 3’ extremity of TAD (B).

